# LSO:Ce Inorganic Scintillators are Biocompatible with Neuronal and Circuit Function

**DOI:** 10.1101/579722

**Authors:** Aundrea F. Bartley, Kavitha Abiraman, Luke T. Stewart, Mohammed Iqbal Hossain, David M Gahan, Abhishek V. Kamath, Mary K. Burdette, Shaida Andrabe, Stephen H. Foulger, Lori L. McMahon, Lynn E. Dobrunz

**Author notes:** These authors contributed equally to the data collection and preparation of this manuscript (Aundrea F. Bartley, Kavitha Abiraman). Corresponding Authors: Lori L. McMahon, PhD Professor, Department of Cell, Developmental, and Integrative Biology, University of Alabama at Birmingham, 1918 University Blvd., MCLM 552, Birmingham AL, 35294; Lynn E. Dobrunz, PhD, Professor, Department of Neurobiology, University of Alabama at Birmingham, 1825 University Blvd, SHEL 902, Birmingham, AL 35294.

## Abstract

Optogenetics is widely used in neuroscience to control neural circuits. However, non-invasive methods for light delivery in brain are needed to avoid physical damage caused by current methods. One potential strategy could employ x-ray activation of radioluminescent particles (RPLs), enabling localized light generation within the brain. RPLs composed of inorganic scintillators can emit light at various wavelengths depending upon composition. Cerium doped lutetium oxyorthosilicate (LSO:Ce), an inorganic scintillator that emits blue light in response to x-ray or UV stimulation, could potentially be used to control neural circuits through activation of channelrhodopsin-2 (ChR2), a light-gated cation channel. Whether inorganic scintillators themselves negatively impact neuronal processes and synaptic function is unknown, and was investigated here using cellular, molecular, and electrophysiological approaches. As proof of principle, we applied UV stimulation to 4 μm LSO:Ce particles during whole-cell recording of CA1 pyramidal cells in acutely prepared hippocampal slices from mice that expressed ChR2 in glutamatergic neurons. We observed an increase in frequency and amplitude of spontaneous excitatory postsynaptic currents (EPSCs), indicating UV activation of ChR2 and excitation of neurons. Importantly, we found that LSO:Ce particles have no effect on survival of primary mouse cortical neurons, even after 24 hours of exposure. In extracellular dendritic field potential recordings, we observed no change in strength of basal glutamatergic transmission up to 3 hours of exposure to LSO:Ce microparticles. However, there was a slight decrease in the frequency of spontaneous EPSCs in whole-cell voltage-clamp recordings from CA1 pyramidal cells, with no change in current amplitudes. No changes in the amplitude or frequency of spontaneous inhibitory postsynaptic currents (IPSCs) were observed. Finally, long term potentiation (LTP), a synaptic modification believed to underlie learning and memory and a robust measure of synaptic integrity, was successfully induced, although the magnitude was slightly reduced. Together, these results show LSO:Ce particles are biocompatible even though there are modest effects on baseline synaptic function and long-term synaptic plasticity. Importantly, we show that light emitted from LSO:Ce particles is able to activate ChR2 and modify synaptic function. Therefore, LSO:Ce inorganic scintillators are potentially viable for use as a new light delivery system for optogenetics.

## Introduction

Over the past decade, the field of optogenetics has expanded our knowledge about the role of individual neuronal cells and specific brain circuits in behavior and disease states (Gradinaru et al., 2009; Yizhar et al., 2011; Lim et al., 2013; Gunaydin et al., 2014; Emiliani et al., 2015; Fenno et al., 2015; Rost et al., 2017; Selimbeyoglu et al., 2017; Barnett et al., 2018). Optogenetics relies on the expression of exogenous light activated ion channels, including the blue light-activated channelrhodopsin-2 (ChR2), that causes depolarization, or the orange-light activated halorhodopsin, which causes hyperpolarization, of the membrane potential of brain cells of interest. Despite these great advances, improvements to the method are constantly being developed (Rein and Deussing, 2012; Lim et al., 2013; Lin et al., 2017; Chen et al., 2018). The use of optogenetics *in vivo* requires delivery of light into the brain, which is most commonly done through implantation of fiber optic waveguides (fibers) or light emitting diodes (LEDs). These can be as large as several hundred microns, which causes damage to delicate brain tissues, especially when implanted deep within brain structures (Aravanis et al., 2007; Ozden et al., 2013; Canales et al., 2018). In addition, attenuation of light occurs through absorption and scattering in brain tissue, resulting in the need for higher intensities of light at the source. This illumination often results in enhanced local temperatures, a consequence that is greater with stronger and more frequent irradiance (Senova et al., 2017), and can lead to as much as a 30% increase in local neuron firing rates (Stujenske et al., 2015). Lastly, glial scarring can occur at the light source (Podgorski and Ranganathan, 2016), decreasing effective light intensities and leading to variability in neuronal control. Currently, there are very few options for noninvasive methods of light delivery into the brain for optogenetics.

Efforts are ongoing to develop and refine minimally invasive strategies for generation of light within brain to combat the challenges mentioned above (Chen et al., 2018). One potential strategy could employ the use of x-rays to activate radioluminescent materials (Shuba, 2014; French et al., 2018). Radioluminescence is the production of visible light by a material exposed to ionizing radiation. Radioluminescent particles (RLPs) can be generated from inorganic scintillator material, which would emit light at different wavelengths depending upon its composition. RLP technology could be superior to current invasive methods because it obviates the need for implanting devices into brain tissue that can cause damage and lose effectiveness over time. In addition, the light will be generated locally and have less attenuation due to tissue scattering. This should allow for much lower power densities needed to achieve opsin activation, thereby reducing nonspecific heat-related effects to the tissue. Finally, the uniformity of light delivery will reduce non-specific effects of a graded response due to light absorption and scatter from a single point source. However, it is unknown whether these inorganic RLPs themselves will impact neuronal processes and synaptic function.

One of the most common inorganic scintillators used in medical imaging is Cerium-doped lutetium oxyorthosilicate (LSO:Ce). LSO:Ce crystals are used as detectors in medical imaging devices because they absorb x-rays and γ-rays extremely well (high density of 7.41 g/cm^3^), have a high light output (~30,000 photons/MeV) and extremely fast decay kinetics (~ 40 ns) (Melcher and Schweitzer, 1991; Roy et al., 2013). The polycrystalline powder form of LSO:Ce has similar properties to the single crystal (Lempicki et al., 2008), and would be within the appropriate size range to be used as a RLP in less invasive optogenetic control of brain circuits. Importantly for optogenetics, activation of LSO:Ce by x-rays emits light in the appropriate range to activate channelrhodopsin-2 (ChR2) (Melcher and Schweitzer, 1991). Since it is a rare-earth metal, it is considered to have a low degree of toxicity. However, the effect of LSO:Ce particles on neurons, which are extremely sensitive to their environment, has not been studied.

Here we demonstrate that LSO:Ce particles, applied for 24 hours, had no effect on neuronal survival in primary cortical cultures. Using adult rat hippocampal brain slices, LSO:Ce particles did not alter basal synaptic excitatory field potentials (fEPSPs) at CA3-CA1 synapses, however, there was a slight decrease in the frequency of spontaneous excitatory post-synaptic currents (sEPSCs) in CA1 pyramidal cells. LSO:Ce particles did not impair the induction of LTP, although there was a slight reduction in LTP amplitude. Importantly, we also demonstrate that light emitted from LSO:Ce microparticles following irradiation is able to activate ChR2 and modulate synaptic function. Together, these results show that LSO:Ce particles are not overtly toxic to neurons or brain slices, although there are modest effects on baseline synaptic function and plasticity. Therefore, the LSO:Ce particles are a potentially viable tool for a less invasive form of in vivo optogenetics.

## Materials and Methods

### 1. Animals

Approval was obtained for all experimental protocols from the University of Alabama at Birmingham Institutional Animal Care and Use Committee. All experiments were conducted in accordance with the Guide for the Care and Use of Laboratory Animals adopted by the National Institutes of Health. Young adult male Sprague Dawley rats were housed in a 22 ± 2° C room with food and water ad libitum. For a subset of experiments, young adult C57Bl/6 mice and adult Emx:ChR2 mice were used. The expression of channelrhodopsin-2 (ChR2) in excitatory neurons was accomplished by crossing Emx-cre mice (B6.129S2-Emx1^tm1(cre)Krj^/J, JAX# 005628) (Gorski et al., 2002) with floxed ChR2/EYFP mice (Ai32 (RCL-ChR2 (H134R)/EYFP, JAX # 012569) (Madisen et al., 2012). C57Bl/6 and Emx:ChR2 mice were housed in a 26 ± 2°C room with food and water ad libitum. Mice were housed with the whole litter until weaned (P25-P28). Weaned mice were housed with no more than 7 mice in a cage. Mouse genotypes were determined from tail biopsies using real time PCR with specific probes designed for cre and EYFP (Transnetyx, Cordova, TN). Pregnant mice (CD-1) were group-housed (4 mice to a cage) in a 22 ± 2° C room with food and water ad libitum and were monitored daily prior to use for neuronal cultures. The animal rooms maintained a standard 12-hour light/dark cycle. 2. Incubation of LSO:Ce microparticles with Acute Hippocampal Slices

### 2. Incubation of LsoO:Ce microparticles with Acute Hippocampal Slices

Acute hippocampal slices were allowed to recover for an hour before application of either vehicle (artificial cerebral spinal fluid, aCSF) or 0.2 to 0.5 mg/mL of LSO:Ce radioluminescent micoparticles. The slices were incubated with and without LSO:Ce microparticles for at least an hour, and up to 6 hours. Incubation of microparticles with hippocampal slices were performed using two types of chambers. Experiments measuring spontaneous postsynaptic currents and baseline synaptic transmission properties used a small incubation chamber that contained 5 mLs of aCSF and was bubbled with 95% O2, pH 7.35–7.45 for the incubation period. For immunohistochemistry and LTP experiments, slices were placed in 2 mLs of aCSF on top of a piece of filter paper within a humidified oxygenated interface recovery chamber, both for recovery and microparticle incubation.

### 3. Electrophysiology using Rats

#### 3.1 Rat Hippocampal Slice Preparation

Young adult male Sprague Dawley rats (age 6–10 weeks; Charles River Laboratories) were anesthetized with isoflurane, decapitated, and brains removed; 400 μm coronal slices from dorsal hippocampus were made on a VT1000P vibratome (Leica Biosystems) in oxygenated (95%O2/5%CO2) ice-cold high sucrose cutting solution (in mm as follows: 85.0 NaCl, 2.5 KCl, 4.0 MgSO_4_, 0.5 CaCl_2_, 1.25 NaH2PO_4_, 25.0 glucose, 75.0 sucrose). After cutting, slices were held at room temperature from 1 to 5 h with continuously oxygenated standard artificial cerebral spinal fluid (aCSF) (in mm as follows: 119.0 NaCl, 2.5 KCl, 1.3 MgSO_4_, 2.5 CaCl_2_, 1.0 NaH_2_PO_4_, 26.0 NaHCO_3_, 11.0 glucose).

#### 3.2 Electrophysiology - Field Recordings

Control and LSO:Ce incubated slices were interleaved to account for slice health. Extracellular field excitatorypostsynaptic potentials (fEPSPs) recorded from the dendritic region in hippocampal area CA1 were performed in a submersion chamber perfused with standard aCSF at room temperature. All data were obtained using the electrophysiology data acquisition software pClamp10 (Molecular Devices, LLC, Sunnyvale, CA.) and analyzed using Clampfit within the pClamp10 suite, and Graphpad Prism 7 (GraphPad Software, Inc.). For CA3-CA1 synapses, Schaffer collateral axons were stimulated using a twisted insulated nichrome wire electrode placed in CA1 stratum radiatum within 200–300 μm of an aCSF-filled glass recording electrode, and paired-pulse facilitation (PPF) characteristic of this synapse (Wu and Saggau, 1994) was recorded. Baseline fEPSPs were obtained by delivering 0.1 Hz stimulation for 200 μs to generate fEPSPs of 0.2-0.3 mV in amplitude. Only experiments with ≤ 10% baseline variance were included in the final data sets.

##### 3.2.1 Input - output curves

After a stable 10 min baseline, input-output (I/O) curves were generated by increasing the stimulus intensity (20 μA increments) until a maximal fEPSP slope was obtained, usually at 200 μA. Initial slope of the five fEPSPs generated at each stimulus intensity were averaged and plotted as a single value. Statistical significance was determined by 2-way ANOVA with Sidak’s multiple comparison test.

##### 3.2.2 Paired Pulse Ratio

After a 10 min stable baseline, pairs of stimulation were delivered at a 50 millisecond (ms) inter-stimulus interval (ISIs). The paired pulse ratio (PPR) was calculated by dividing the initial slope of the second fEPSP by the initial slope of the first fEPSP.

##### 3.2.3 Long Term Potentiation

At CA3-CA1 synapses, following a 10 min stable baseline (0.1 Hz, 200 μs with stimulation intensity set to elicit initial fEPSP amplitude of 0.3-0.4 mV), NMDA receptor (NMDAR)-dependent LTP was induced using high-frequency stimulation (HFS, 100 Hz, 1 s duration × 5, 60 s interval). Statistical significance was determined using unpaired Student’s *t*-test by comparing the average of the fEPSP slope from the last 5 min of the recording (35– 40 min) to baseline for each genotype (*p < 0.05).

#### 3.3 Electrophysiology - Whole Cell

All recordings were performed in a submersion chamber with continuous perfusion of oxygenated standard ACSF. Whole-cell voltage clamp recordings were carried out in blind patched CA1 pyramidal neurons. IPSCs were pharmacologically isolated with bath perfusion of DNQX (10uM; Sigma) and DL-AP5 (50uM; Tocris). Spontaneous IPSCs were recorded using CsCl internal solution (in mM: 140.0 CsCl, 10.0 EGTA, 5.0 MgCl_2_, 2.0 Na-ATP, 0.3 Na-GTP, 10.0 HEPES; E_C1_ = 0mV). All cells were dialized for 3-7min prior to the beginning of experimental recordings. Stability of series resistance during the recording was verified posthoc through comparing the average rise and decay time of sIPSCs. Recordings were discarded if the rise or decay time changed by > 20%.

### 4. Electrophysiology in Mice

#### 4.1 Mouse Hippocampal Slice Preparation

Male and female mice, 4 to 7 months of age, were anesthetized with isoflurane and sacrificed by decapitation using a rodent guillotine. The brains were rapidly removed and placed in ice cold dissection solution containing the following (in mM): 135 N-Methyl-D-glucamine, 1.5 KCl, 1.5 KH_2_PO_4_, 0. 5 CaCl2, 3.5 MgCl_2_, 23 NaHCO3, 0.4 L-Ascorbic acid, and 10 glucose, bubbled with 95% O2/5% CO2, pH 7.35–7.45, and osmolarity 295–305 (Albertson et al., 2017). A vibratome (Campden 7000smz-2, Lafayette Instrument) was used to cut 300 μM thick hippocampal brain slices. The slices were maintained for 45–60 min at 37°C in oxygenated recovery solution containing (in mM) 120 NaCl, 3.5 KCl, 0.7 CaCl2, 4.0 MgCl_2_, 1.25 NaH_2_PO_4_, 26 NaHCO_3_, and 10 glucose and then kept at room temperature. Slices were stored at room temperature in a holding chamber containing the recovery solution and bubbled with 95% O2/5% CO2 for >30 minutes before recording.

#### 4.2 Whole Cell Electrophysiology Recording

Photostimulation of the microparticles occurred by using pulses of UV light (315 nm or 365 nm, 500 ms pulse duration). The 365 nm UV light was generated by a Colibri.2 LED Light Source (Zeiss) applied to acute hippocampal slices through a 10x objective, resulting in an illumination area of 5.29 mm^2^. The 365 nm light intensity was measured as being approximately 0.003 mW/mm^2^ by ThorLabs PM100D Optical Power Meter. The 315 nm UV light was generated by a custom built portable optogenetic unit using components obtained from ThorLabs. The 315 nm UV light is applied to the acute hippocampal slice through a 2.5 mm stainless steel ferrulle, therefore allowing a similar illumination area to that of the 10x objective. Both the Colibri.2 and the custom built 315 nm portable unit were triggered by a Master-9 digital stimulator (A.M.P.I).

For the recordings, slices were placed in a submersion recording chamber and perfused (3-4 ml/min) with aCSF. Experiments were performed at 28-32°C. For spontaneous EPSC (sEPSC) recordings, CA1 pyramidal cells were blindly patched on a Zeiss Examiner A1 upright microscope. Neurons were patched in the voltage-clamp configuration and recorded at a holding potential of −60 mV using an Axopatch 200B amplifier (Molecular Devices). Patch electrodes (4–6 MΩ) were filled with a potassium gluconate based internal solution composed of the following (in mM): 130 K-gluconate, 0.1 EGTA, 20 KCl, 2 MgSO_4_·7H_2_O, 10 HEPES, 5 phosphocreatinetris, 10 ATP, and 0.3 GTP. The pH was adjusted to 7.2 with KOH and osmolarity was 290-295. The access resistance and holding current (<200 pA) were monitored continuously. Recordings were rejected if either access resistance or holding current increased ≥ 25% during the experiment.

Analysis of sEPSC frequency and amplitude were performed using custom software written in Visual Basic, which measured amplitude and interevent interval. Events were fit to a template response and all events that fit the template and passed visual inspection were included in the analysis.

### 5. Immunohistochemistry

Acute 300 μm hippocampal slices were obtained from P50 to P70 C57Bl/6 mice. Slices were incubated with and without 0.25 to 0.5 mg/mL of LSO:Ce radioluminescent particles for 0, 1, 3, and 6 hours. Slices were fixed overnight at 4°C in 4% paraformaldehyde in 0.1 M phosphate buffer saline (PBS) and immunohistochemistry was performed based on a modified protocol (Dissing-Olesen and MacVicar, 2015; Miller et al., 2017; Yang et al., 2018). Briefly, the free-floating fixed slices were washed 5 times for 10 minutes in PBS. The slices were permeabilized for 1 hour at room temperature in 0.1 M PBS (pH 7.4) with 0.3% Triton X-100 and 20% DMSO (PBSTD). Slices were blocked with 10% donkey serum (cat# 017-000-121, Jackson ImmunoResearch Inc.) in PBSTD for 3 hours at room temperature. Following blocking, the sections were immunostained with a monoclonal mouse antibody against GFAP (1:1000, cat# 3670, Cell Signaling Technology) for 48 to 96 hours at 4°C in blocking solution. After washing, the slices underwent another blocking step. Then the slices were incubated with Alexa Fluor 488 conjugated polyclonal donkey anti-mouse antibody secondary (cat# 715-545-150, Jackson ImmunoResearch Inc.) diluted 1:400 in blocking solution for 16 to 40 hours at 4°C. Staining specificity was confirmed by omission of primary antibody. Sections were mounted with the nuclear dye DAPI (4’, 6-diamindino-2-phenylindole dihydrochloride) in the VECTASHIELD HardSet mounting medium (cat# H-1500, Vector Laboratories). Immunostained slices were imaged using an epifluorescent Nikon eclipse ni microscope and NIS Elements v. 4.20.02 software. Analysis of images was performed using ImageJ (Fiji).

### 6. Primary cortical neuronal culture

Mouse primary cortical neuronal culture from pregnant CD-1 mice was made as previously described (Andrabi et al., 2014). Embryos obtained from mice at Day 15–16 of gestation were used to prepare primary cortical neuron culture. Briefly, the cortical regions of the embryonic brains were aseptically dissected, freed of meninges and dissociated in dissecting medium (DMEM + 20% FBS) and subjected to trypsin digestion at 37°C for 5 min. Tryptic digestion was stopped by the addition of dissecting medium and the cell suspension were centrifuged at 1000 g for 5 min. Next, the pelleted cells were subjected to mechanical trituration in complete Neurobasal medium (10 mM glucose, 1 mM GlutaMAX-I, 1 mM Na-Pyruvate and 2% B-27) and passed through a 40 μm filter. The cells were plated to a density of 5 ×10^5^ cells/ml. On day in vitro (DIV) 1, the cultures were treated with 5-fluoro 2-deoxyuridine (40 μM) to inhibit glial cell growth and proliferation. Experiments were performed between DIV 10 and 11. Under these conditions, mature neurons represent 90% of the cells in the culture.

### 7. Cell death assay

Neurons at DIV 10 were incubated with either microparticles (LSO:Ce, dispersed in PBS), at concentrations ranging from 0.05 to 0.2 mg/ml or PBS for 24 hours. Cell viability after addition of nanoparticles was determined by using Alamar blue reagent, a water-soluble resazurin dye (blue colored) which is reduced to red fluorescent resorufin dye by metabolically active cells. Alamar blue reagent [10% (v/v)] was then added to each well containing Neurobasal medium and incubated for 3 hours at 37°C. Blank control well containing microparticles only were used to exclude possible interactions with the assay. Neurobasal medium containing Alamar blue reagent was added to these wells. Following 3 hours of incubation, 100 μL of the medium was collected from each well and transferred to a 96-well microplate. The fluorescence was measured at the excitation and emission wavelength of 540 and 595 nm, respectively using a microplate reader. The fluorescence values were normalized by the control (PBS) and expressed as percent viability.

### 8. Materials

Commercial lutetium oxyorthosilicate (LSO:Ce) particles (median particle size: 4 μm) were purchased from Phosphor Technologies and were doped with cerium at a 1-10 atomic % cerium. Prior to their use, the particles were washed with deionized water and vacuum air dried.

## Results

### Light emitted from LSO:Ce particles can weakly activate channelrhodopsin-2

LSO:Ce particles emit within the activation spectrum for channelrhodopsin-2 (ChR2). Interestingly, these particles can be activated with ultraviolet (UV) light. As a proof of principle, we took advantage of this property to test whether light emitted from LSO:Ce microparticles could enhance synaptic activity. However, it was necessary to first test whether ChR2 is activated by UV light. We used EmxCre:ChR2 mice to determine the extent to which ChR2 is activated by various wavelengths of UV light. Using UV light at 365 nm, we observed a photocurrent (−22.4 ± −2.3 pA, n=10,) in recordings from CA1 pyramidal cells in acute slices, even with a relatively low light intensity (0.003 mW/mm^2^). We were concerned that the activation of ChR2 with 365 nm light would hamper our ability to see activation of ChR2 from light generated by the LSO:Ce particles. Therefore, we built a portable UV system that would emit light at 315 nm. Application of 315 nm light onto CA1 pyramidal cells that express ChR2 generated an extremely small, but detectable, photocurrent (−5.5 ± −2. 0 pA, n=5).

As the light emitted from the LSO:Ce particles might not be uniform enough to generate a synchronized synaptic event, we analyzed the impact on the amplitude and frequency of sEPSCs. As there is cell to cell variability with sEPSCs, we used a within cell control (no UV light) to normalize the amplitude and frequency of sEPSCs. Even though only a small photocurrent (−5.5 ± 2.0 pA, n=5) was generated with application of 315 nm light, we observed an increase in frequency of sEPSCs during UV light exposure (Figure 1A,D). Importantly, the enhancement is much larger when the LSO:Ce particles are present (Figure 1B, D). A similar enhancement of the frequency of sEPSCs was seen when 365 nm UV light was applied (UV only 149.4 ± 20.9 % vs UV + LSO:Ce 225.2 ± 24.4 %, n = 8, 7, Student’s t-test, *p* = 0.04). Interestingly, the amplitude of the sEPSCs are decreased with the application of 315 nm light (Figure 1C), which might be an indication that the sEPSCs are shifted to more action potential independent events. However, the addition of the LSO:Ce particles enhances the amplitude of the sEPSCs compared to UV alone (Figure 1C). Additionally, the overall size of photocurrent induced by the 315 nm UV light was enhanced in the presence of the particles as compared to UV light application alone (Vehicle slices −5.5 ± −2.0 pA, vs LSO:Ce slices −12.3 ± 1.8 pA, n=5,7, Student’s t-test, *p* = 0.03). These data indicate that the light emitted from LSO:Ce particles is able to activate ChR2 and increase the frequency of sEPSCs.

**Figure 1.**
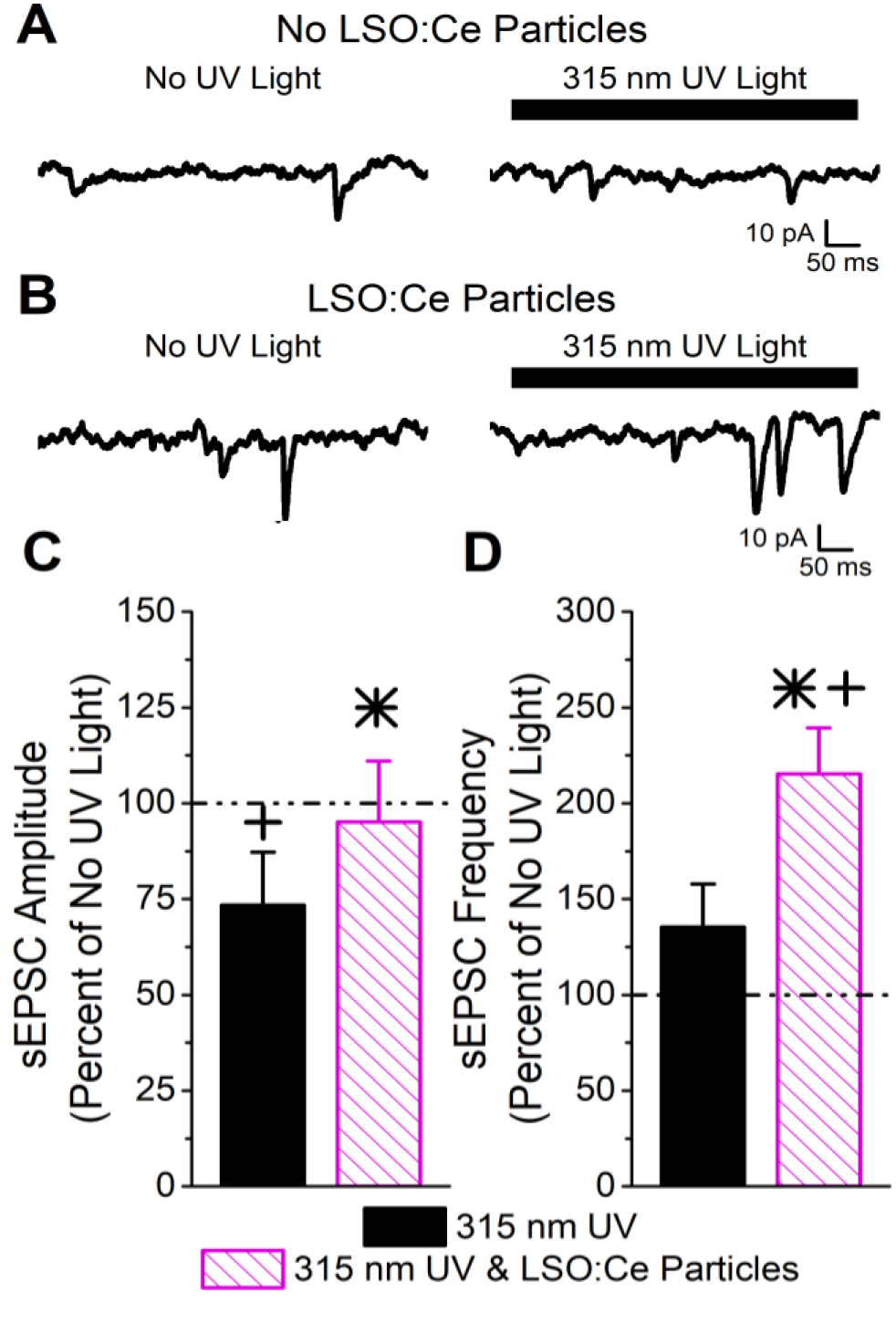
Light emitted from LSO:Ce microparticles enhances synaptic transmission from ChR2 expressing CA1 pyramidal cells. A) Example traces of sEPSCs recorded from a CA1 pyramidal cell in an acute hippocampal slice from a Emx:ChR2 mouse that was not incubated with LSO:Ce particles. The solid line above the traces represents the section that was analyzed, either in the presence or absence of 500 ms of 315 nm UV light. B) Example traces of sEPSCs onto CA1 pyramidal cells from an acute hippocampal Emx:ChR2 mouse slice that was incubated with LSO:Ce particles for 1 hour. The solid line above the traces represents the section that was analyzed, either in the presence or absence of 500 ms of 315 nm UV light. C,D) For each experiment, the sEPSC amplitude and frequency measured in the presence of the 315 nm UV light was normalized to a section of the same trace that was not exposed to UV light to account for cell to cell variability. C) Application of UV light alone decreased the amplitude of sEPSCs compared to baseline (n=5, Student’s t-test, *p* = 0.01). Nevertheless, the presence of the LSO:Ce particles enhanced the sEPSC amplitude as compared to UV alone (n = 5, 7, Student’s t-test, *p* = 0.03). D) There was a trend for exposure to UV light alone to enhance the frequency of sEPSCs (n=5, Student’s t-test, *p* = 0.19). The light emitted from LSO:Ce particles by UV activation almost doubles the number of sEPSCs compared to baseline (n = 5, 7, Student’s t-test, *p* = 0.04). * indicates significant difference (*p*< 0.05) with and without LSO:Ce particles + indicates significant difference (*p*< 0.05) compared to no UV light

### Prolonged exposure to LSO:Ce microparticles does not alter neuronal survival or properties

We next tested the biocompatibility of LSO:Ce microparticles. Neurons are extremely sensitive to changes in their environment and any perturbations could lead to neuronal cell death. We determined if prolonged exposure to LSO:Ce particles had any effect on neuronal survival using the Alamar Blue assay and primary cortical cultures. The result demonstrates no significant sign of toxicity exerted by nanoparticles at any of the doses tested, even with 24 hours of exposure to the LSO:Ce microparticles (Figure 2).

**Figure 2.**
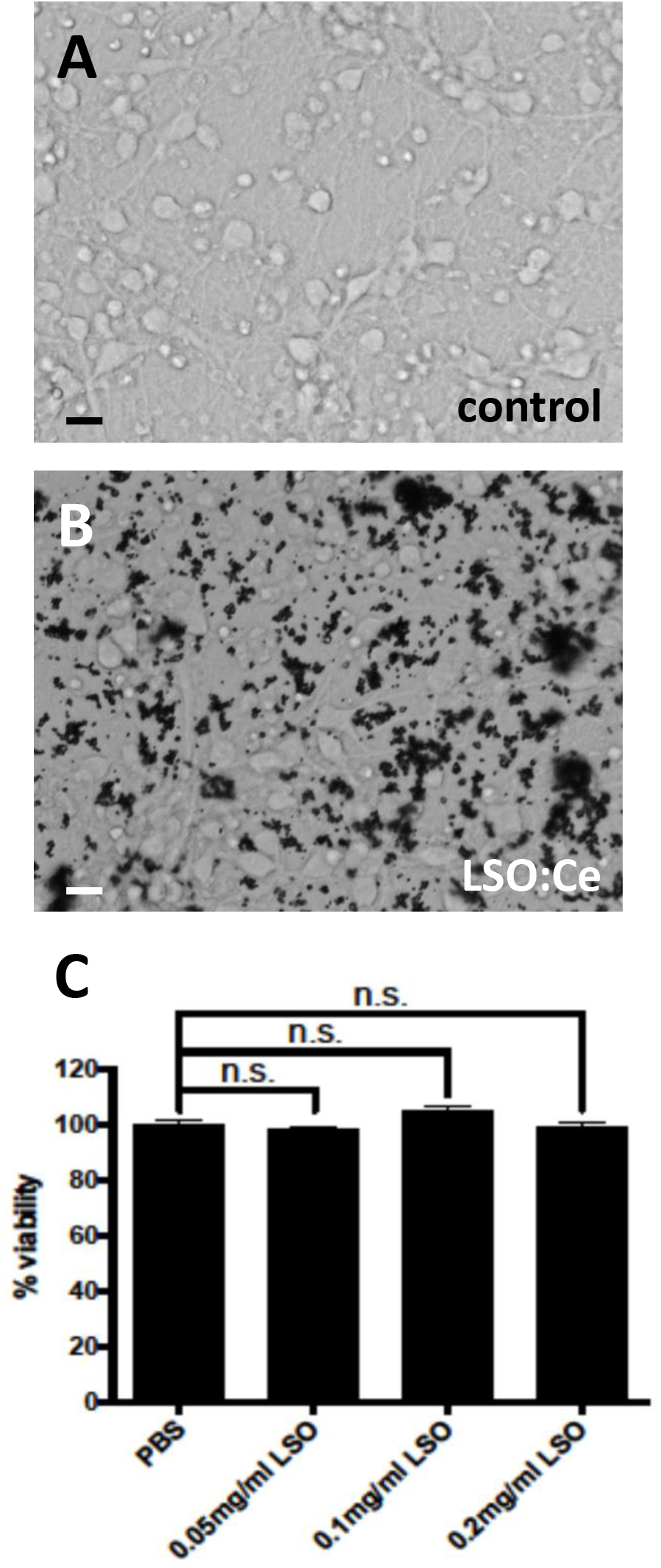
Prolonged exposure of LSO:Ce microparticles is not toxic to neurons. A, Representative image of primary cortical neurons 11 DIV B, Representative image of primary cortical culture 11DIV incubated in LSO:Ce particles for 24 hours. C, Group data at indicated LSO:Ce concentrations were added to neurons at 10 DIV and cell death assay was performed on DIV 11 using Alamar Blue reagent. Results are expressed as percent of control. Data represent as mean ± SEM, n = 6, One-way ANOVA with Tukey’s post hoc test. n.s, not significant

Primary cortical cultures are more sensitive to extracellular perturbations than acute slices. However, we confirmed the results that neuronal survival was not affected by acute application of the LSO:Ce microparticles using acute hippocampal slices. We incubated slices with 0.25-0.5 mg/mL of LSO:Ce for 0-6 hours, and manually counted the number of DAPI positive neurons in the pyramidal layer (Figure 3). We saw no difference in the density of CA1 pyramidal cells between unexposed and exposed slices even after 6 hours of treatment (Vehicle slices 109.0 ± 8.9 cells/mm^2^ vs LSO:Ce slices 105.9 ± 16.3 cells/mm^2^, n = 11, 9, Student’s t-test, *p* = 0.86). Additionally, acute incubation of LSO:Ce particles did not change the resting membrane potential of CA1 pyramidal cells (Vehicle slices −63.9 ± 2.7 mV vs LSO:Ce slices −66.1 ± 2.1 mV, n = 16, 13, Student’s t-test, *p* = 0.55). However, the input resistance of the cells was slightly reduced (Vehicle slices 99.1 ± 6.6 MΩ vs LSO:Ce slices 80.4 ± 5.1 MΩ, n = 16, 13, Student’s t-test, *p* = 0.04). Overall, exposure to LSO:Ce did not affect neuronal health.

**Figure 3.**
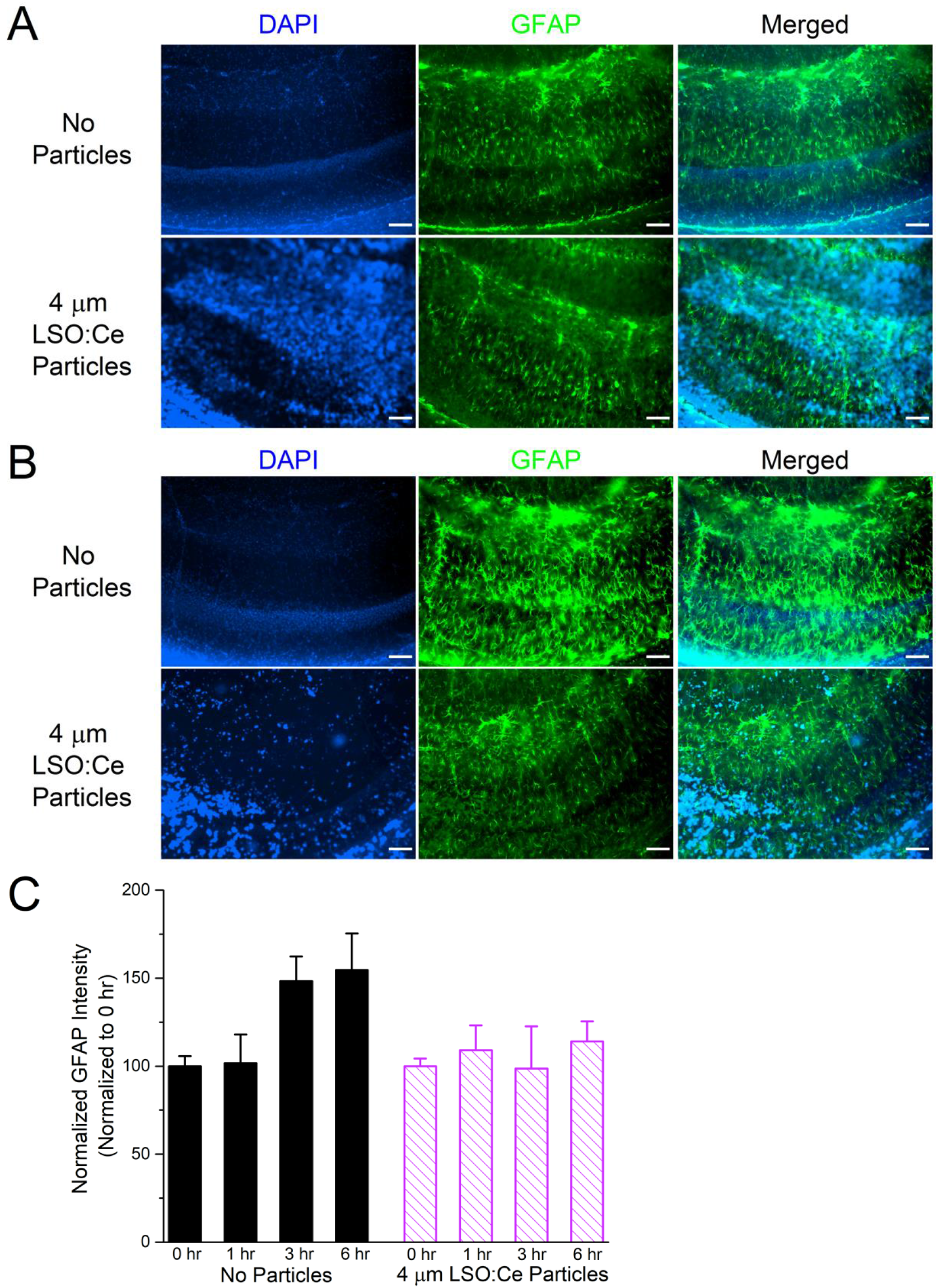
Exposure of LSO:Ce microparticles to acute hippocampal slices has does not elevate GFAP. A) Example images of GFAP staining in acute hippocampal slices immediately after application of vehicle or LSO:Ce particles. Merged images include DAPI (blue) and GFAP (green) staining. Scale bars: 100 μm B) Example images of GFAP staining in acute hippocampal slices 6 hours after application of vehicle or LSO:Ce particles. Merged images include DAPI (blue) and GFAP (green) staining. Scale bars: 100 μm C) Group data shows that the intensity of GFAP staining is not altered with the addition of the LSO:Ce particles (n = 8 slices/4 animals per condition, ANOVA, F_(3,63)_ = 1.76, *p* = 0.17)

As implanted probes have been shown to increase glial scarring (Canales et al., 2018), we wanted to determine if astrocytes were modified in the presence of these particles. We used immunohistochemistry to analyze glial fibrillary acidic protein (GFAP) in LSO:Ce preincubated slices compared to control slices. As the LSO:Ce particles could be activated at the wavelengths necessary to visualize GFAP staining, we performed control experiments in which we immediately processed slices after application of vehicle (aCSF) and the particles (Figure 3) to test for differences in fluorescent intensity caused by light being emitted or absorbed by the particles. However, the overall fluorescent intensity of GFAP staining was not different between vehicle-treated slices and LSO:Ce-treated slices at the 0 hour time point, indicating that measurement of GFAP fluorescence was not altered due to the emission and absorbance capabilities of the LSO:Ce particles (Vehicle slices 1.03 × 10^6^ ± 1.14 × 10^5^ vs LSO:Ce slices 1.02 × 10^6^ ± 6.02 × 10^4^, n = 8, 8, Student’s t-test, *p* = 0.98). Slices were incubated for up to 6 hours with either vehicle or the LSO:Ce microparticles. The results show that there was no increase in the level of GFAP staining due to prolonged incubation with the LSO:Ce particles (Figure 3).

### LSO:Ce microparticles modestly alter synaptic transmission

Even though overall cell health was unaltered by exposure to LSO:Ce particles, synaptic transmission could still be affected. We further analyzed this by incubating hippocampal slices with LSO:Ce particles for 1-3 hours, and followed by recording extracellular dendritic fields potentials (fEPSPS) from CA1 pyramidal cells in stratum radiatum in response to increasing the number of CA3 Schaffer collaterals stimulated (Figure 4A). The initial slope of the fEPSP represents the postsynaptic response, and we found no difference in the maximal response generated between the control and LSO:Ce-treated slices (Figure 4B). However, we saw a small, but significant, increase in the paired-pulse ratio (PPR), suggesting that the LSO:Ce particles may alter presynaptic function (Figure 4B).

**Figure 4.**
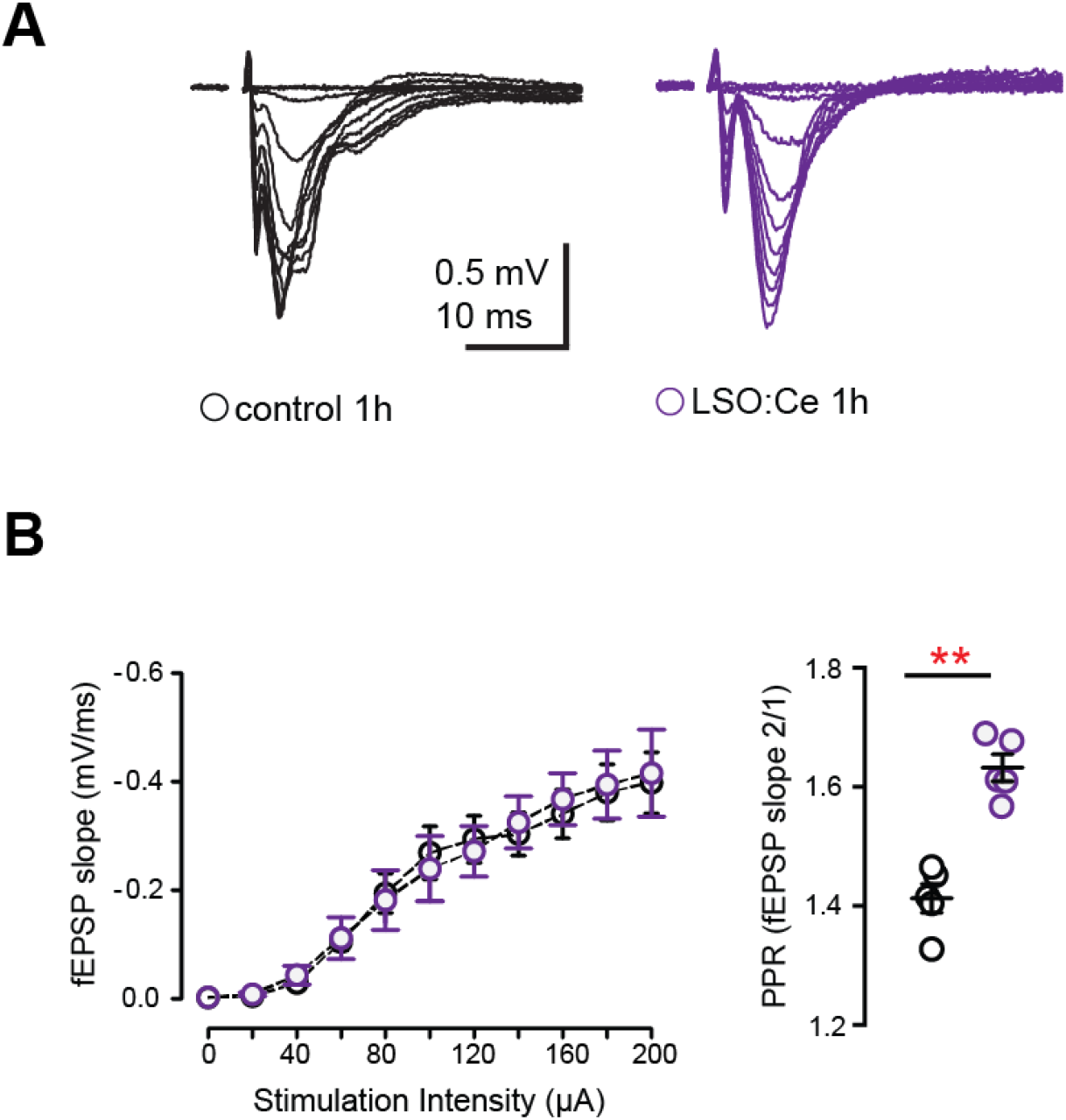
Acute application of LSO:Ce microparticles has no effect on the strength of basal synaptic transmission at CA3-CA1 synapses. A) Representative traces of field excitatory postsynaptic potentials (fEPSPs) at CA3-CA1 synapses in response to increasing stimulus intensity from acute hippocampal slices incubated without and with LSO:Ce particles (black traces: control and purple traces: particle). B) No change in the input/output (I/O) curve in in the presence of LSO:Ce particles (n = 6 slices/6 animals control, n = 6 slices/6 animals LSO:Ce; *p > 0.05). Data represent mean ± SEM. Significance determined by 2-way ANOVA with Sidak’s multiple comparison test. Paired-pulse ratio (PPR) was significantly increased following incubation with particles (n = 6 slices/6 animals control, n = 6 slices/6 animals LSO:Ce; *p <0.05)

To further investigate whether incubation with LSO:Ce particles affects synaptic transmission, we preincubated acute hippocampal slices with LSO:Ce particles for 1 to 3 hours. We then performed whole cell recordings from CA1 pyramidal cells and recorded spontaneous excitatory (sEPSCs) (Figure 5) and inhibitory postsynaptic currents (sIPSCs) (Figure 6). There was no change in the amplitude of sEPSCs (Figure 5), however the frequency of the events was slightly but significantly reduced with exposure to LSO:Ce particles. On the other hand, sIPSCs were unaltered in the presence of the particles (Figure 6). This suggests either that the particles are reducing the number of excitatory synapses, altering axonal excitability, or modifying presynaptic function.

**Figure 5.**
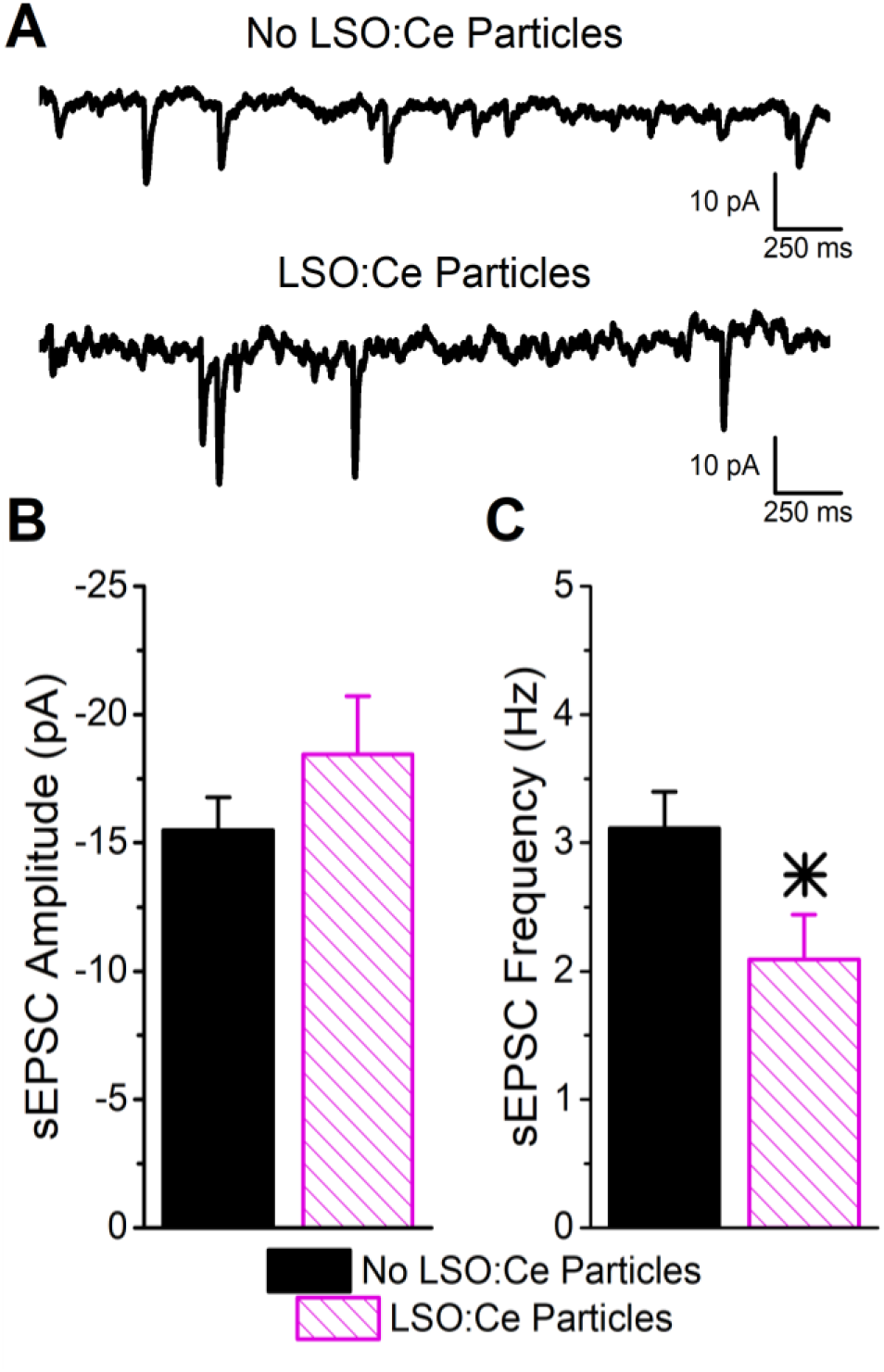
Acute application of LSO:Ce microparticles reduces the frequency, but not amplitude, of sEPSCs recorded from CA1 pyramidal cells. A) Example traces of spontaneous EPSCs onto CA1 pyramidal cells from acute hippocampal slices incubated with and without the particles for 1 hour. B) Group data showing that the amplitude of sEPSCs was unaltered in the presence of LSO:Ce particles (n = 15, 11, Student’s t-test, *p* = 0.24). C) Group data showing the frequency of sEPSCs was significantly reduced in the presence of LSO:Ce particles (n = 15, 11, Student’s t-test, *p* = 0.03). * indicates significant difference (*p*< 0.05) with and without LSO:Ce particles

**Figure 6.**
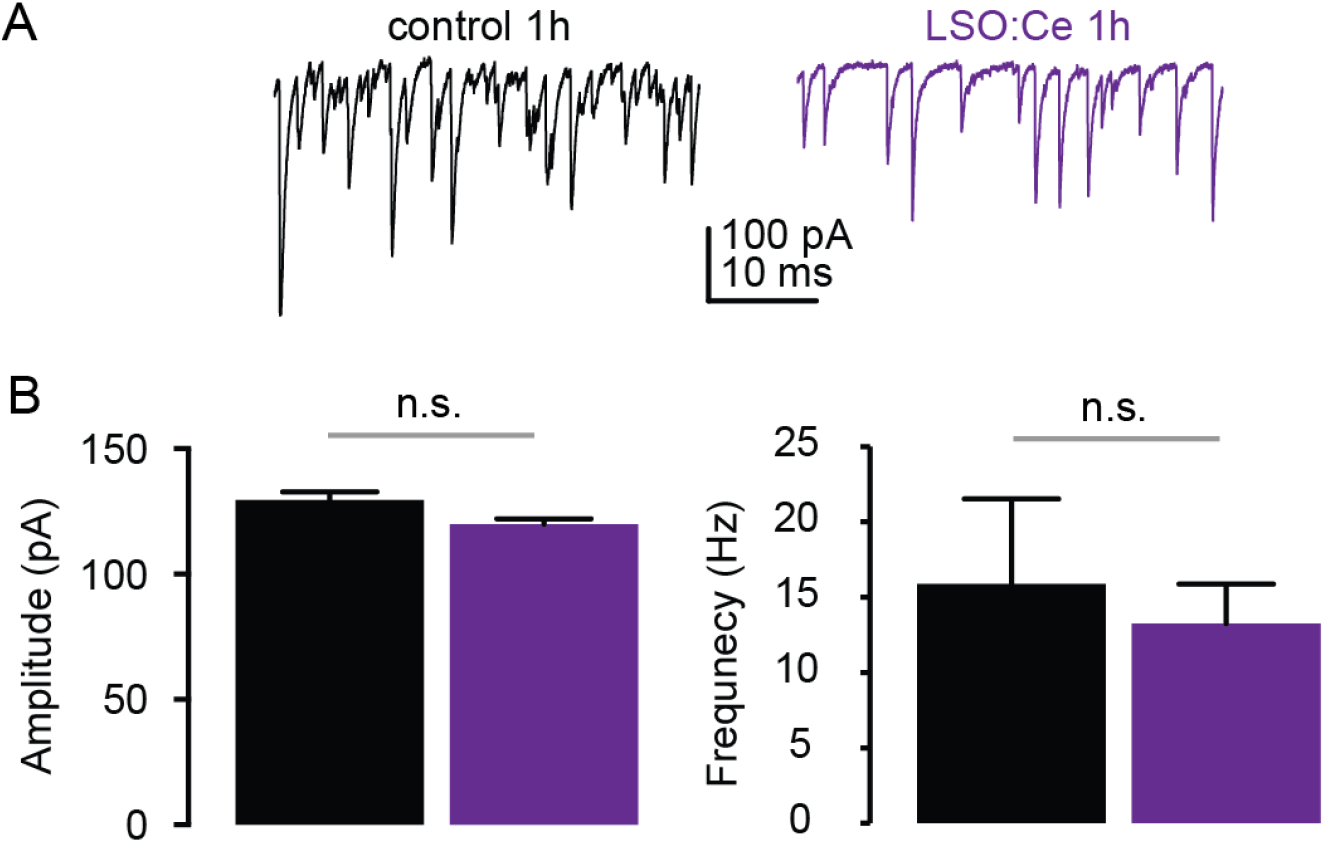
Acute application of LSO:Ce microparticles had no effect on sIPSCs recorded from CA1 pyramidal cells. A) Example traces of sIPSCs onto CA1 pyramidal cells from acute hippocampal slices incubated with and without the particles for 1 hour. B) Group data showing the amplitude of sIPSCs was unaltered in the presence of LSO:Ce particles (n = 5, 9, Mann-Whitney test, *p* = 0.18). C) Group data showing that frequency of the sIPSCs was unaltered in the presence of LSO:Ce particles (n = 5, 9, Mann-Whitney test, *p* =0.79).

### LSO:Ce microparticles modestly alter Long Term Potentiation

Next, we asked if long-term plasticity, a more robust measure of synaptic health and integrity, is compromised by pre-incubation with LSO:Ce particles. Long term potentiation (LTP), an increase in synaptic strength that lasts for at least 40 min *in vitro* and hours to days *in vivo* and is thought to underlie learning and memory, has been extensively characterized at CA3-CA1 synapses, and involves insertion of excitatory AMPA receptors (AMPARs) on post-synaptic CA1 pyramidal cells. We again incubated acute hippocampal slices for 1-3 hours with LSO:Ce particles, and then recorded extracellular fEPSPs in response to Schaffer collateral stimulation in an interleaved fashion where control and LSO-Ce treated slices were alternated. Following establishment of a 10-minute baseline, we electrically induced LTP at CA3-CA1 synapses by stimulating Schaffer collaterals at 100 Hz for 0.5 seconds, 5 times, separated by intervals of 20 seconds. While this protocol produced identical post-tetanic potentiation and LTP up to 20 min post-tetanus in control and LSO:Ce treated slices, there was a very small but significant decrease in LTP expression at 40 min post-tetanus in the LSO:Ce treated versus control slices, which may suggest that the particles have a slightly negative impact on long-term expression of LTP (Figure 7).

**Figure 7.**
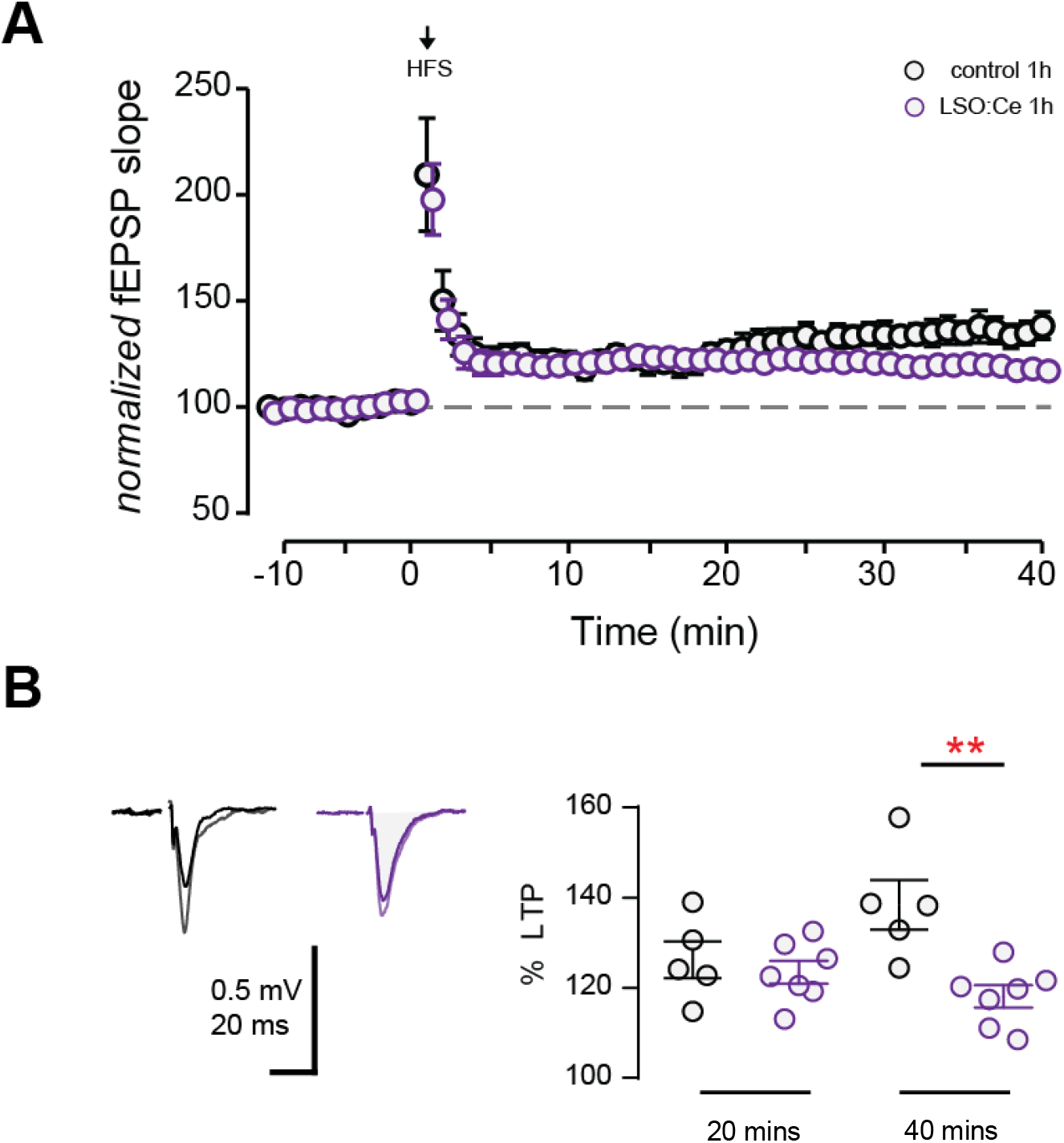
Acute application of LSO:Ce microparticles decreases LTP at CA3-CA1 synapses. A) The magnitude of High Frequency Stimulation (HFS, 5X 1 sec at 100Hz) induced LTP at CA3-CA1 synapses was significantly decreased in slices incubated with particles. B) Averaged representative responses at −5 and +35 minutes HFS (black and grey traces: control and purple and light purple traces: particle). Deficit in late but not early LTP magnitude in slices incubated without and with particles for an hour (40 min LTP: 138±5% and 118±2%, n = 5 slices/5 animals, n = 7 slices/7 animals respectively; *p = 0.0038; 20 min LTP: 124±4% and 121±2%, n = 5 slices/5 animals, n = 7 slices/7 animals respectively; *p >0.05). Data represent mean ± SEM. Significance determined by unpaired Student’s *t*-test.

## Discussion

Our results provide proof of principle that light from LSO:Ce microparticles can activate ChR2 and modulate synaptic function. In addition, we demonstrate that LSO:Ce is not overtly toxic to neurons, as there was no effect of the particles on neuronal survival in culture even with incubations as long as 24 hours. Synaptic function and plasticity also appear to be intact, as there was no change in the input/output relationship in CA1, and LTP was able to be induced. However, there were indications that LSO:Ce particles themselves have effects on synaptic function and plasticity, although the effects were small. Together, these results suggest that LSO:Ce particles could potentially be suitable for *in vivo* optogenetics.

LSO:Ce was introduced as a scintillator in the early 90s (Melcher and Schweitzer, 1992) and has been extensively characterized since. Cerium forms the luminescence centers in LSO, and the Ce doped LSO displays a high emission intensity under X-ray illumination, with an emission maximum at 420 nm. This emission peak makes LSO:Ce well suited to optogenetics applications using ChR2, which involves activation with 473 nm light. The use of X-rays has the distinct advantage over LED to activate ChR2, as X-rays are able to penetrate the skull, removing the need for an invasive delivery method.

Implantation of LEDs or a fiber requires the light output to be proportionally higher than necessary to activate ChR2 because of light scattering properties in brain tissue. One study using a fiber to deliver light reported that local tissue temperature increased by 0.8 °C (Senova et al., 2017). Interestingly in that study, no effect was seen on neuronal cell death with acute light applications at the highest illumination used (Senova et al., 2017), however the question still remains how the tissue can handle local heating with longer duration or repeated applications.

If the conditions are right, tissue heating under illumination can cause damage and contribute to observed behavioral or physiological effects (Long and Fee, 2008). The use of nanoparticles made from inorganic scintillators will allow for the light source to be extremely close to ChR2 and therefore less light would be needed to activate ChR2, reducing the risk of damage caused by heat.

Here we observed that light emitted from LSO:Ce microparticles in response to UV stimulation caused an increase in the frequency of sEPSCs onto CA1 pyramidal cells, indicating activation of ChR2. However, we did not detect firing of action potentials or large light-evoked EPSCs, suggesting that the activation of ChR2 is modest. UV light alone is also able to provide weak activation of ChR2, as seen by small photocurrents and an increase in sEPSC frequency. The photocurrents were modest with stimulation at 365 nm, and even smaller at 315 nm. One limitation of the current experiment is that the particles were applied to the surface of the slice, and therefore are likely to be 100 microns or more from the cell expressing ChR2. The proposed use of RLPs for *in vivo* optogenetics would use smaller particles that will be located much closer to ChR2, thus reducing light attenuation from scattering. As a result, the light needed for ChR2 activation *in vivo* should be even less.

We observed no toxicity in primary neuronal cultures following 24 hours of LSO:Ce microparticle exposure, and in acute hippocampal slices, there was no inflammatory response assessed by GFAP staining in astrocytes or decrease in strength of basal transmission assessed by I/O curves at CA3-CA1 synapses. However, there were some minor effects on synaptic function. While there was no effect on sIPSCs, there was a small reduction in the frequency but not amplitude of sEPSCs together with an increase in PPR of evoked excitatory transmission, indicating an effect of the particles on presynaptic release probability (Dobrunz and Stevens, 1997). There was also a small reduction in the input resistance of CA1 pyramidal cells, which could contribute to reduced cell excitability. Alterations in neuronal excitability have been shown to modulate presynaptic release probability (Crabtree et al., 2017), suggesting that this could be the underlying mechanism for the minor effects on synaptic function with application of the LSO:Ce microparticles.

We also observed a slight but significant decrease in the magnitude of LTP, the long-term enhancement of synaptic strength and a cellular correlate of learning and memory (Malenka and Bear, 2004). Because LTP induction and expression is highly susceptible to cellular and behavioral stress, it is a sensitive measure of neuronal and synaptic health. LTP was induced using high frequency stimulation and was monitored for 40 min post-tetanus. While the LSO:Ce particles did not affect the magnitude of the post-tetanic potentiation or the magnitude of LTP up to 20 mins post-tetanus, there was a slight but significant decrease in the LTP magnitude at 40 mins post-tetanus in the LSO:Ce treated slices. This suggests the possibility of a small effect on the long-term expression of LTP, however whether this change is biologically relevant to learning and memory is unknown.

Overall, the effects on synaptic and intrinsic neuronal properties were modest, indicating that the LSO:Ce particles do not majorly disrupt synaptic transmission and circuit function. Whether these minor effects on synaptic transmission are a result of physical interaction and the weight of the particles on the delicate neuronal structures is not clear. However, if toxic chemicals were leaching out of the particles to negatively impact cell health, we would likely have observed cell death in the primary neuronal cultures after 24 hours of exposure. Future studies are needed to determine if there are detrimental effects of LSO-Ce particles with UV or X-ray exposure potentially through the release of free radicals, as observed with Cd quantum dots (Jamieson et al., 2007). However, any potential toxic effects could be overcome through PEGylation of the particles (Cheng et al., 2011).

Together, our results support the possibility of using LSO-Ce particles combined with ChR2 expression to non-invasively regulate synaptic circuit function *in vivo*.

## Abbreviations

aCSF-: artificial cerebral spinal fluid
ChR2-: channelrhodopsin-2
fEPSP–: field excitatory postsynaptic potentials
GFAP-: glial fibrillary acidic protein
LED-: light emitting diodes
LSO:Ce–: Cerium doped lutetium oxyorthosilicate
LTP–: long term potentiation
PBS-: phosphate buffer saline
RLPs-: radioluminescent particle
sEPSCs–: spontaneous excitatory postsynaptic currents
sIPSCs–: spontaneous inhibitory postsynaptic currents
UV-: ultraviolet

## Acknowledgments

This work was funded by NSF Track 2 FEC OIA-1632881 to SHF.

## Author Contributions Statement

LD, LM AB, KA, LS, and SA designed the experiments, AB, KA, LS, DG, AK, and MH performed experiments and analyzed the data, LD, LM AB, KA, SF, MB, MH, and SA wrote and edited the manuscript.

